# BRANEart: identify stability strength and weakness regions in membrane proteins

**DOI:** 10.1101/2021.08.22.457277

**Authors:** Sankar Basu, Simon S. Assaf, Fabian Teheux, Marianne Rooman, Fabrizio Pucci

## Abstract

Understanding the role of stability strengths and weaknesses in proteins is a key objective for rationalizing their dynamical and functional properties such as conformational changes, catalytic activity, and protein-protein and protein-ligand interactions. We present BRANEart, a new, fast and accurate method to evaluate the per-residue contributions to the overall stability of membrane proteins. It is based on an extended set of recently introduced statistical potentials derived from membrane protein structures, which better describe the stability properties of this class of proteins than standard potentials derived from globular proteins. We defined a per-residue membrane propensity index from combinations of these potentials, which can be used to identify residues which strongly contribute to the stability of the transmembrane region or which would, on the contrary, be more stable in extramembrane regions, or *vice versa*. Large-scale application to membrane and globular proteins sets and application to tests cases show excellent agreement with experimental data. BRANEart thus appears as a useful instrument to analyze in detail the overall stability properties of a target membrane protein, to position it relative to the lipid bilayer, and to rationally modify its biophysical characteristics and function. BRANEart can be freely accessed from http://babylone.3bio.ulb.ac.be/BRANEart.

## 1 Introduction

The broad family of integral membrane proteins provides indispensable components of living cells. Being embedded in biological lipid membranes, these proteins include important scaffolds and functional sites, which bind targeted molecules floating around in the cytosol or in the extracellular medium. They therefore serve as attractive drug targets [1, 2].

Stability and physico-chemical characteristics greatly vary between transmembrane (TM) and extramembrane (EM) regions of an integral membrane protein due to the difference in their surrounding chemical environment. EM domains in the cytosolic or extracellular medium resemble globular proteins as their surfaces are exposed to water, while residues embedded within the membrane are exposed to lipids and are thus characterized by an elevated hydrophobicity [3, 4]. Because of this mixed environment, the study of folding and stability of membrane proteins is very challenging and only few tools have been dedicated to investigate them [5, 6, 7, 8]. Another reason why membrane proteins are less studied than globular proteins is their lower number of experimental three-dimensional (3D) structures [9].

Just as for globular proteins, the native structure of a membrane protein corresponds to the global minimum of the free energy landscape in physiological conditions. In general, however, some protein residues or regions, taken individually, are not in their global free energy minimum and correspond to what we call stability weaknesses [10, 11, 12]. Indeed, not all residues can simultaneously adopt their lowest free energy conformations, because of the polypeptide chain which constraints the relative motions of the residues. Native conformations can be viewed as the best compromise between conflicting interactions. We define strong and weak residues as residues that are, or are not, optimized for stability properties, respectively. For example, stability weaknesses occur when a contact between two residues in the native structure does not have a favorable free energy contribution and can thus easily break, change conformation, and/or remain flexible. They often play a key role in protein function, interactions, and conformational changes. The concept of stability weaknesses is close to the notion of frustration except that the latter also includes kinetic constraints on fast folding [13].

To further advance these issues, we implemented and expanded a series of new statistical mean-force potentials designed to specifically describe the stability properties of membrane proteins, which were first defined in [3]. Based on these potentials, here we derived a per-residue membrane propensity index which predicts whether residues situated in the membrane have a stabilizing contribution or would prefer to be in EM regions, and similarly for residues outside the membrane. This index thus identifies strong and weak residues in EM and TM regions, and can also be used to predict how a protein is, or is not, inserted in the membrane.

It has to be noted that we are studying in this paper relative, EM/TM, strengths and weaknesses, defined from the difference in folding free energy according to whether a residue is in one or the other region. They are different from the strengths and weaknesses defined in globular proteins as residues of which the folding free energy contribution is either highly optimal or not optimal at all, as for example computed by the SWOTein webserver [12].

We made BRANEart freely available in the form of a web server called BRANEart, designed to help the scientific community to explore stability strengths and weaknesses in membrane proteins, which is a key element in the study of their stability and function.

## 2 MATERIALS AND METHODS

### 2.1 Protein structure data sets

The membrane protein data set 𝒟_mem_ comes directly from a recent study [3]. It consists of 163 X-ray structures of integral membrane proteins that have been collected from the Protein Data Bank (PDB) [14]. The selected structures have a resolution of at most 2.5 Å and a pairwise sequence identity of at most 30%, computed using the PISCES protein sequence culling server [15].

The set 𝒟_mem_ contains 107 *α*-helical and 52 *β*-barrel integral proteins, and 4 *α*-helical bitopic integral proteins that do not span the lipid bilayer completely. Each of these proteins was annotated using OPM (Orientations of Proteins in Membranes) [16], a curated web resource that positions the biological lipid bilayer on experimentally resolved structures of integral membrane proteins and membrane-bound peptides. Using these annotations, the TM and EM portions of each protein were further segregated in two subdata sets 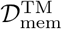 and 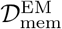.

All selected protein chains in 𝒟_mem_ were considered in the context of their biological assembly (or biounit), which corresponds to their functional quaternary conformation. This ensures a more realistic representation of the surrounding protein environment experienced by the protein. The biological units were taken to be those defined by the authors of the X-ray structures or, in absence of annotations from the authors, as those predicted by the Protein Interfaces, Surfaces and Assemblies (PISA) tool [17].

Finally, we also set up a second data set 𝒟_glob_ of 4,860 monomeric globular protein structures from the PDB, to be used as an independent set for validating our method. The proteins from this data set have a monomeric biological unit, a good quality X-ray structure with a maximum resolution of 2.5 Å and a pairwise sequence identity of at most 25%.

The list of all proteins belonging to the data sets 𝒟_mem_ and 𝒟_glob_ are given in the GitHub repository github.com/3BioCompBio/BRANEart.

### 2.2 Statistical potentials

Statistical potentials are coarse-grained mean-force energy functions derived from frequencies of associations of sequence and structure motifs in a data set of known protein structures. These frequencies are transformed into free energies using the inverse Boltzmann law [18, 19, 20]. These potentials depend on the characteristics of the data set from which they are derived. For example, temperature-dependent statistical potentials are obtained from protein data sets of different melting temperatures [21, 22], and solubility-dependent potentials from proteins of different solubility [23].

Here we derived a series of membrane protein potentials from the sets 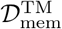 and 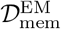, extending the work of [3]. More precisely, we considered the following first and second order statistical potential terms [20]:

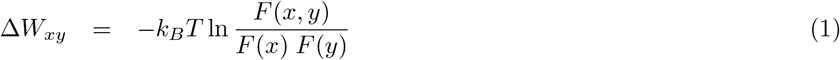

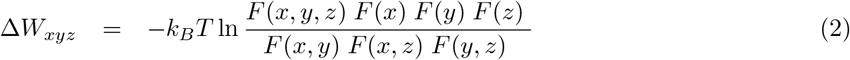

where *k*_*B*_ is the Boltzmann constant, *T*, the absolute temperature conventionally taken to be room temperature, and *F*, the relative frequencies computed from a given data set of protein structures. The variables *x, y, z* stand for any of the four elementary structure or sequence descriptors *s, d, t* and *a*: *s* is an amino acid type, *d*, the spatial distance between the average side chain geometric centers of two residues separated by at least one residue along the polypeptide chain [19], *t*, a (*ϕ, ψ, ω*) backbone torsion angle domain [24], and *a*, a solvent accessibility bin where the solvent accessibility is defined as the ratio (in %) between the solvent accessible surface area of a residue in the structure and in an extended Gly-X-Gly conformation [19].

We constructed two versions of each of the potentials defined by Eqs (1)-(2). In the first, all frequencies *F* were computed from the structure set 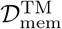, *i*.*e*. considering only protein regions embedded in the lipid membrane. In the second, all frequencies *F* were computed from 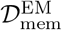, thus considering only extramembrane protein regions. The potentials extracted from TM regions, 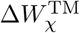 with *χ* = *xy* or *xyz*, describe the stability properties of membrane proteins inside the lipid membrane, while the potentials extracted from EM regions, 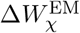, describe the stability properties outside the lipid bilayer. Note that the only membrane protein potentials that we constructed and analyzed earlier are the inter-residue distance potentials 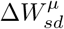 and 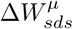 [3], where *µ* is either EM or TM.

The full list of 19 membrane statistical potentials derived here and their characteristics are given in Section 1 of Supplementary Materials.

### 2.3 Per-residue folding free energies

The contribution of each residue *i* in a protein to its overall folding free energy computed with a given potential 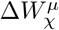, noted 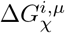, was computed in such a way that an equal amount was allocated to each of the residues that carry the structural descriptors *a, t* and *d* included in *χ*. According to the number of residues that the structural descriptors encompass, we used the following rules [11, 12]:

- if the structure descriptors involved in *χ* are localized on a single residue *i* (which is the case when e.g., *χ* = *st, sa, sst, ssa*), the total contribution is assigned to that residue;
- if the structure descriptor is localized on two residues *i* and *j* (e.g., *χ* = *sd, tt, aa, sds, saa, stt*) half of the energy contribution is assigned to each of these residues;

For the potential *χ* = *ss* that does not contain any structure descriptor, an equal amount was allocated to each residue carrying a sequence descriptor *s*.

For example, let us consider the three potentials 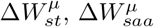 and 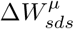. The per-residue folding free energy contributions are:

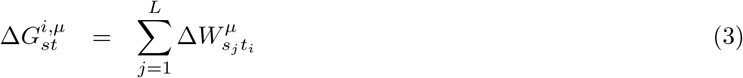

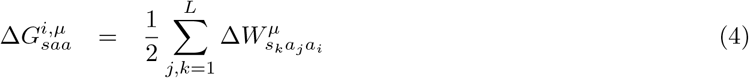

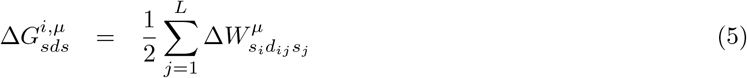

where *i, j* and *k* denote positions along the amino acid sequence and *L* the sequence length. For further details, we refer the reader to previous studies [11, 3, 12].

In a last step, the folding free energy values so obtained for each residue *i* were smoothed by taking a weighted average over the 5-residue sequence window [*i* − 2, *i* + 2] [3]:

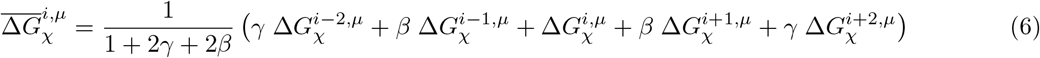

The weighting parameters *γ* and *β* were chosen to minimize the level of weaknesses in the transmembrane regions (see Section 2.5). For the N- and C-terminal residues, the smoothing was done on the residues present in the [*i* − 2, *i* + 2] sequence interval.

In this way, each residue *i* was tagged with two folding free energy values 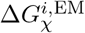 and 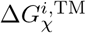 for each statistical potential *χ*, irrespective of its location, in either EM or TM regions.

### 2.4 Membrane propensity Index

We introduced a per-residue membrane propensity index MPr^*i*^ to predict to what extent a residue *i* in a folded protein corresponds to a stability strength or to a weakness when placed in a given, lipid or aqueous, environment. From a physico-chemical perspective, MPr^*i*^ estimates whether a residue shows a preference for the EM or TM environments, and can be interpreted as an index of stability of a residue within its structural context. It is defined as a linear combination of all folding free energy terms derived in the previous subsection:

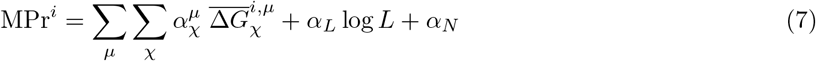

where 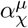 are real-valued parameters that need to be optimized (see Section 2.5). The MPr^*i*^ score is built in such a way that it returns a value close to one for residues predicted to be stable in lipid environments, and a value close to zero for residues predicted to be stable in aqueous environments.

Residues that strongly contribute to the stability of the region to which they belong are considered as stability strengths, and residues that would be more stable elsewhere in the protein are called stability weaknesses.

### 2.5 Training and optimization

To estimate the membrane propensity index MPr^*i*^ introduced in Eq. (7), we have to identify the 40 free parameters introduced in Eqs (6-7). We performed this parameter identification by minimizing the overall amount of structural weaknesses in the proteins from the 𝒟_mem_ set. The general idea behind this procedure comes from the minimal frustration principle [13] stating that proteins have evolved, and still evolve, to optimize the folding energy landscapes.

To perform the parameter identification on the 𝒟_mem_ dataset, we need to know if each of the residues in the proteins of this set is in an EM or TM region. For this, we searched the residue annotations in the OPM database [16]. These annotations were then assigned to the vector 𝒪, with the convention that zero corresponds to an EM region and one to a TM region. 𝒪 served as target function in the model training. Note that the OPM annotations are actually predictions and may also suffer from inaccuracies.

We can now define the cost function 𝒞 as the square of the difference between the OPM annotations and the predicted membrane propensity MPr defined in Eq. (7):

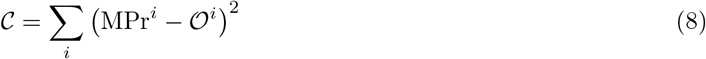

where the sum is over all residues in the training set 𝒟_mem_. The parameter identification was done with Python (v.3) using the regression subroutine LinearRegression. Note that logistic regressions were also tested in the training process, but appeared to yield less good results.

The performance of the predictor was evaluated on the training data set 𝒟_mem_ using a leave-one-out cross validation procedure, in which 𝒟_mem_ but one protein was used as a training set and the removed protein as a test set. The rounds of training and testing were repeated as often as there are proteins in 𝒟_mem_ (163). As a final result of the training, the BRANEart model outputted a continuous membrane propensity score MPr varying approximately between {-0.5, 1.5}. Values that are close to one identify residues that are stable in the phospholipid bilayer, while values close to zero represent residues that are stable in the aqueous solvent.

Note that all the cross validations that we performed in this paper are strict: the target protein is excluded from all steps of our computations, from the derivation of the statistical potentials to the computation of MPr and the identification of *ϕ*_0_, and the sequence identity between learning and test set proteins is at most 30%.

### 2.6 EM/TM prediction

The membrane propensity score MPr was also used to set up a binary classifier that predicts whether residues in a membrane protein belong to TM or to EM regions. For that purpose, we transformed the continuous MPr scores into a discrete binary function, where 0 means EM and 1 TM, using an appropriate cutoff value *ϕ*_0_ such that a residue *i* is predicted to be in the TM region if MPr^*i*^ ≥ *ϕ*_0_ and in the EM region otherwise. The value of *ϕ*_0_ was identified to minimize the difference with OPM assignments.

To evaluate the performance of this classifier, we used the balanced accuracy (BACC) defined as:

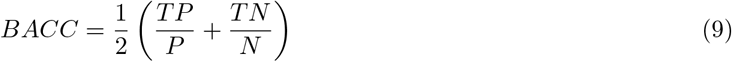

where TP and TN mean true positive and true negatives, respectively, and P an N positives and negatives.

## 3 Results and Discussion

### 3.1 Setting up BRANEart

We set up the BRANEart predictor, which predicts for each residue in a protein structure a membrane protein index MPr, defined as a linear combination of several types of statistical potentials, derived from either EM or TM regions of a set of membrane protein structures (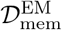 and 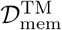), as defined in Eqs (1)-(7). The coefficients of the linear combination were identified to follow OPM annotations [16], through the minimization of Eq. (8), with the definition that an MPr close to one means a preference to be located in a TM region and an MPr close to zero, to be in an EM region. This identification was performed in strict cross validation (Section 2.5). In what follows, when computing MPr on the sets 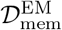 and 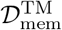, we used the cross-validated values.

The MPr index yields a quantitative measure of the stabilizing or destabilizing contribution that a residue has in a specific environment, in other words, whether it acts as a weakness or a strength in that environment. Note that we compared here aqueous and lipid environments, but that this approach can be generalized to the comparison between other environments.

### 3.2 Large-scale application of BRANEart

We applied BRANEart to the membrane protein data set 𝒟_mem_ and to the globular monomeric protein set 𝒟_glob_ defined in Section 2.1, and computed the per-residue MPr score for the 56,715 and 1,258,648 amino acid residues contained in these two sets. Note that the huge difference in size between these two sets, of more than an order of magnitude, is due to the paucity of experimental structures available for membrane proteins.

As seen in Table 1, the mean ⟨MPr⟩ value is close to zero in globular proteins, ⟨MPr⟩ = 0.13, with a low standard deviation of *σ* = 0.13. This means that the large majority of the residues are stabler in an aqueous than in a lipid environment, which is obviously the case. Residues in membrane proteins have intermediate preferences (⟨MPr⟩ = 0.40) with a much larger *σ* of 0.37. However, splitting membrane proteins into TM and EM regions clarifies these findings: residues in EM regions clearly prefer to be in an aqueous environment (⟨MPr⟩ = 0.18), while residues inside the membrane prefer a lipid environment (⟨MPr⟩ = 0.74). These excellent results are a first validation of BRANEart.

**Table 1:** Mean per-residue index ⟨MPr⟩ and standard deviation *σ* of the distributions computed for the four protein structure sets considered in this paper; *N* is the number of residues in the data set.

| Data set | Protein type | $\langle \text{MPr} \rangle$ | $\sigma$ | $N$ |
| --- | --- | --- | --- | --- |
| $\mathcal{D}_{\text{glob}}$ | Globular proteins | 0.13 | 0.13 | $1.3 \times 10^6$ |
| $\mathcal{D}_{\text{mem}}$ | Membrane proteins | 0.40 | 0.37 | $5.7 \times 10^5$ |
| $\mathcal{D}_{\text{mem}}^{\text{TM}}$ | Trans-membrane regions | 0.74 | 0.27 | $2.3 \times 10^5$ |
| $\mathcal{D}_{\text{mem}}^{\text{EM}}$ | Extra-membrane regions | 0.18 | 0.22 | $3.4 \times 10^5$ |

The MPr distributions are plotted in Fig. 1 for each of the four data sets. They all show a unimodal bell-shaped distribution, except the one computed from 𝒟_mem_ which is more spread out with two peaks, a narrow peak and a flatter one. This distribution is the union of the two unimodal distributions computed from the sets 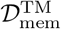 and 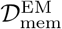, which are shifted relative to each other and show a marked preference for lipid and aqueous environments, respectively. The distribution from 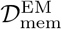 is roughly centered around the same MPr value as the one obtained from 𝒟_glob_, but it is much less peaked although the proteins from both sets are in an aqueous environment. This suggests that there are more stability weaknesses in membrane proteins than in globular proteins or more precisely, that there are more residues in the EM regions of membrane proteins than in globular proteins which would prefer to be in a lipid environment.

**Figure 1:**
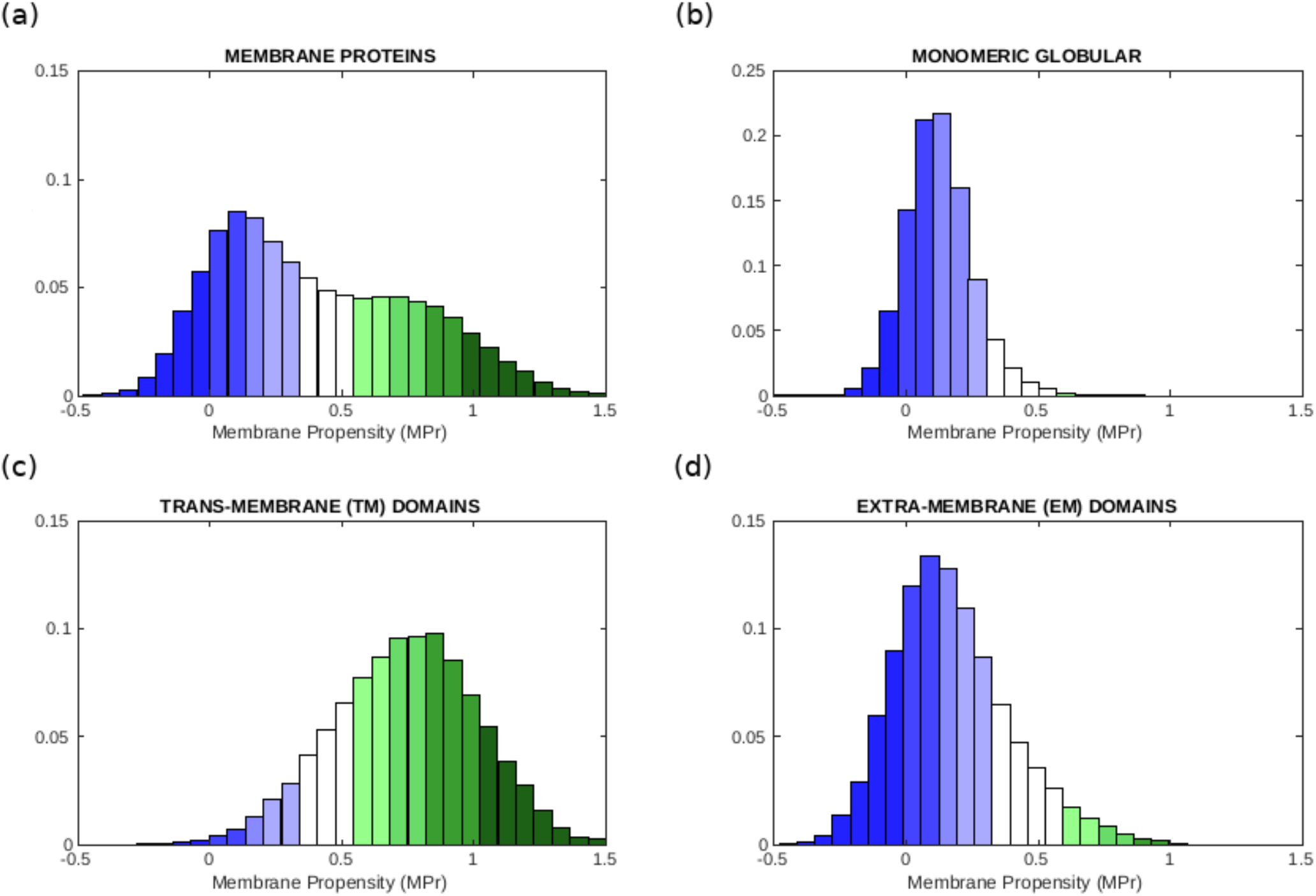
Distribution of MPr scores computed from different of protein data sets: (a) membrane proteins 𝒟_mem_, (b) globular proteins 𝒟_glob_, (c) TM regions 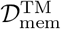 and (d) EM regions 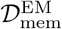. The color code is defined in Table 2.

To graphically visualize here and in the BRANEart webserver the level of stability of a residue, we defined stability classes in terms of the mean and standard deviations of the MPr distribution and we colored the residues accordingly, as explained in Table 2. Residues that are strong in aqueous environments but weak in lipid environments are colored in blue. Residues that are strong in the phosholipid bilayer and weak in aqueous solvent are colored in green. Different color graduations were defined to further differentiate between highly stable, stable, moderately stable and mildly stable residues in EM regions, and equivalently in TM regions.

**Table 2:**
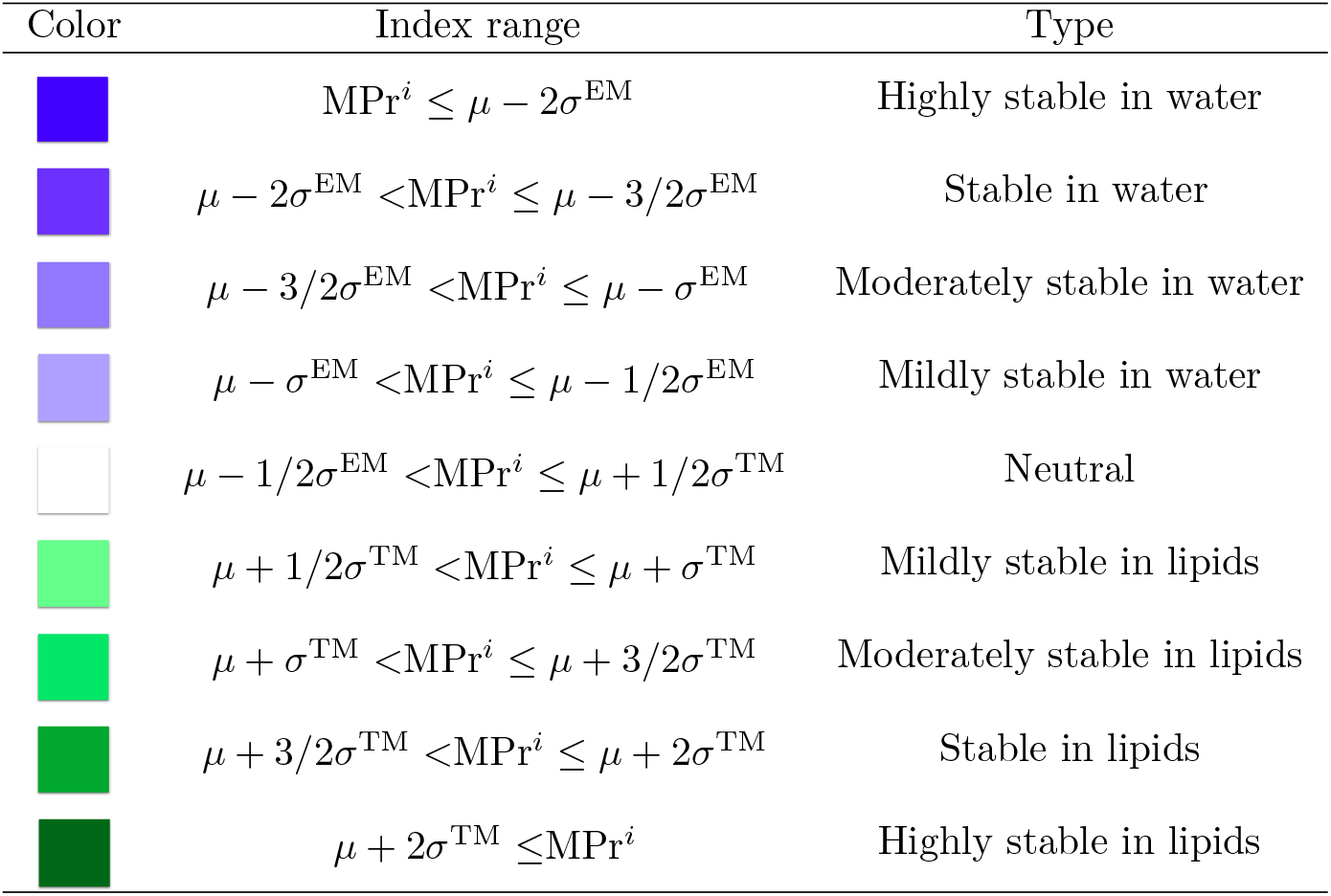
Classes of MPr scores defined in terms of the mean and standard deviations of the MPr distribution. *µ* is the mean of the MPr distribution computed on 𝒟_mem_ and *σ*™ and *σ*^EM^ the standard deviations of the MPr distributions computed from 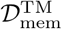 and 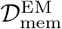, respectively. The colors used in the visualization frameworks of the BRANEart webserver are here defined.

| Color | Index range | Type |
| --- | --- | --- |
| | $\text{MPr}^i \leq \mu - 2\sigma^{\text{EM}}$ | Highly stable in water |
| | $\mu - 2\sigma^{\text{EM}} < \text{MPr}^i \leq \mu - 3/2\sigma^{\text{EM}}$ | Stable in water |
| | $\mu - 3/2\sigma^{\text{EM}} < \text{MPr}^i \leq \mu - \sigma^{\text{EM}}$ | Moderately stable in water |
| | $\mu - \sigma^{\text{EM}} < \text{MPr}^i \leq \mu - 1/2\sigma^{\text{EM}}$ | Mildly stable in water |
| | $\mu - 1/2\sigma^{\text{EM}} < \text{MPr}^i \leq \mu + 1/2\sigma^{\text{TM}}$ | Neutral |
| | $\mu + 1/2\sigma^{\text{TM}} < \text{MPr}^i \leq \mu + \sigma^{\text{TM}}$ | Mildly stable in lipids |
| | $\mu + \sigma^{\text{TM}} < \text{MPr}^i \leq \mu + 3/2\sigma^{\text{TM}}$ | Moderately stable in lipids |
| | $\mu + 3/2\sigma^{\text{TM}} < \text{MPr}^i \leq \mu + 2\sigma^{\text{TM}}$ | Stable in lipids |
| | $\mu + 2\sigma^{\text{TM}} \leq \text{MPr}^i$ | Highly stable in lipids |

Interestingly, the amount of strengths and weaknesses in TM and EM regions are almost identical, as computed from the MPr distributions: 7% of the residues are weak in both regions, 75% are strong and the remaining ones are neutral. Without surprise, most residues thus contribute to the stabilization of the regions to which they belong; note that the functional residues are usually among the weak residues.

We also examined how the MPr values change as a function of the solvent accessibility in globular proteins. Therefore, we divided all the residues in 𝒟_glob_ into three groups: core residues with accessibility ≤ 20%, intermediate residues with 20% *<* accessibility ≤ 50%, and surface residues with accessibility *>* 50%. The average MPr score in these groups gradually drops from core to surface: 0.15 (*σ*=0.15), 0.13 (*σ*=0.10), 0.08 (*σ*=0.10). As seen in Fig. 2, the whole MPr distribution is shifted towards lower values, and becomes more peaked. This reflects the fact that residues in the core of globular proteins locally feel a hydrophobic environment and have a slightly higher MPr, while surface residues are in direct contact with water and a lower MPr.

**Figure 2:**
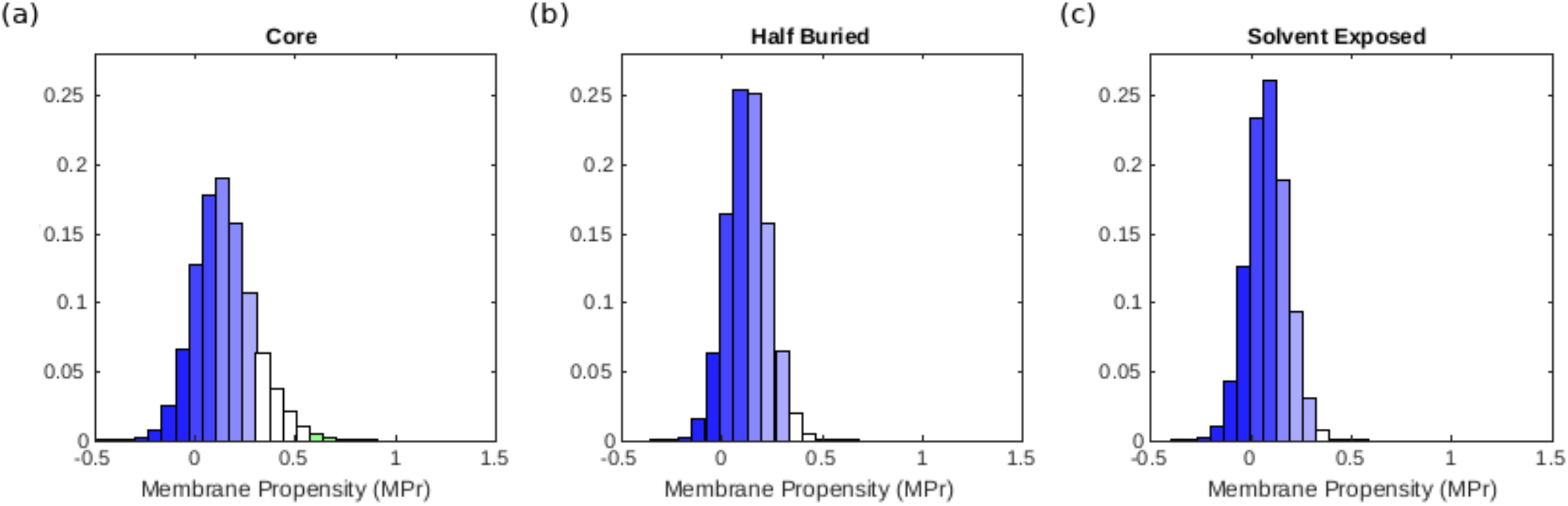
Distribution of MPr scores computed from the globular protein set 𝒟_glob_ for residues that are (a) situated in the core, (b) half buried and (c) solvent exposed. The color code is defined in Table 2.

### 3.3 Application to *α*-helical and *β*-barrel membrane proteins

Membrane proteins of which the TM region has an *α*-helical or a *β*-barrel conformation exhibit different folding and stability properties [3, 4]. Indeed, the former type of proteins are essentially localized in the cytoplasmic membranes of eukaryotic and prokaryotic cells and quite rarely in outer membranes, whereas the latter proteins are found in outer membranes of gram-negative bacteria, mitochondria or chloroplasts.

We observe from the distributions depicted in Fig. 3.a-b that residues pertaining to *α*-helical folds have a bigger preference for lipid environments than those pertaining to *β*-barrel folds. Indeed, average ⟨MPr⟩ values are equal to 0.44 (*σ*=0.40) and 0.34 (*σ*=0.31) for *α* and *β* proteins, respectively. If we focus on the TM regions, this tendency is even more marked (Fig. 3.c-d): ⟨MPr⟩= 0.78 (*σ*=0.28) and 0.63 (*σ*=0.23).

**Figure 3:**
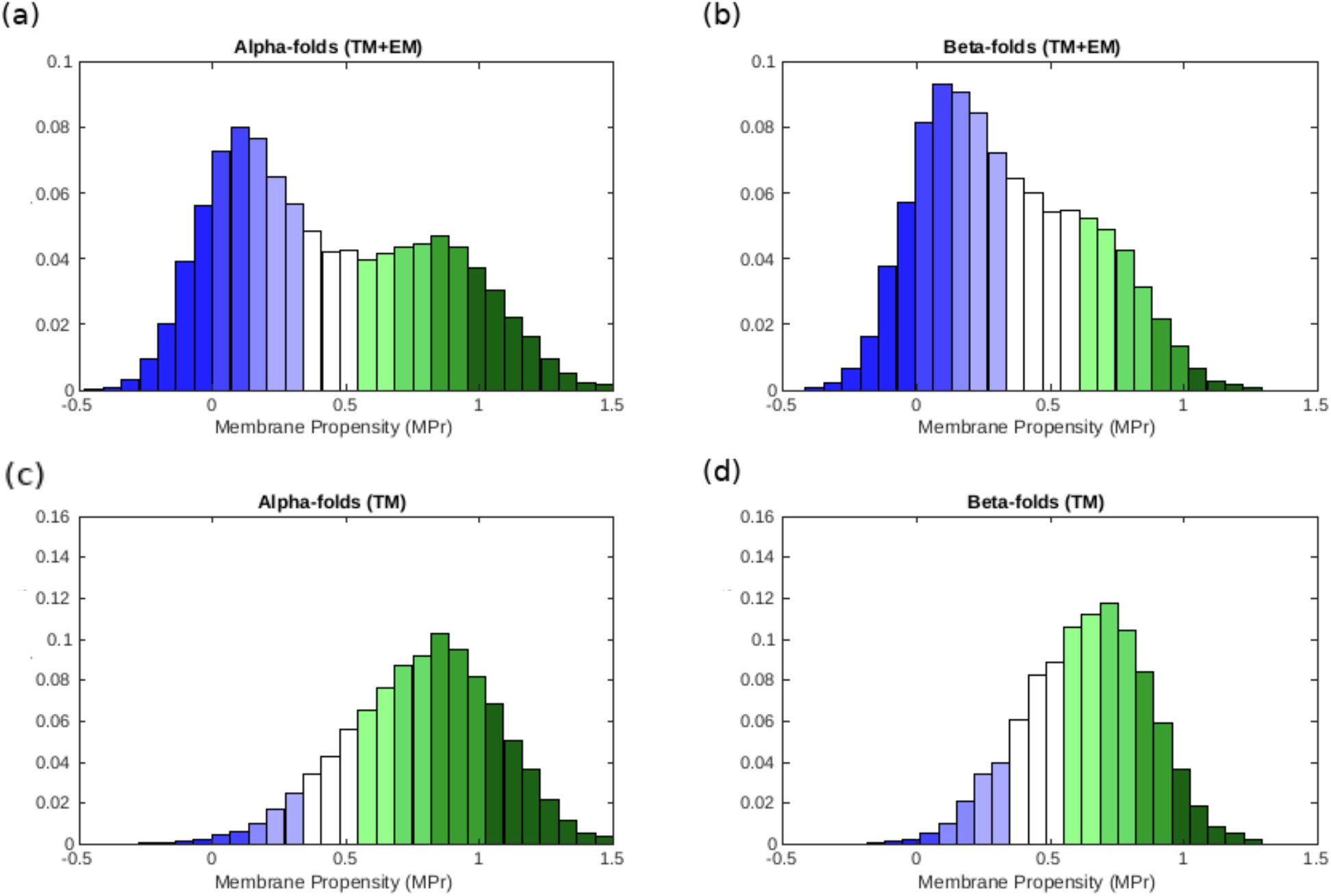
MPr distribution computed from the membrane proteins in 𝒟_mem_ that have an *α*-helical or *β*-barrel structure. (a) *α*-folds (TM and EM), (b) *β*-folds (TM and EM), (c) TM region of *α*-folds and (d) TM region of *β*-folds. The color code is defined in Table 2

These results are in line with a series of facts. First, *α*-helices are coiled structures stabilized by regularly spaced hydrogen bonds between residues at positions *i* and *i* + 4 along the polypeptide chain. In contrast, *β*-sheets are stabilized by hydrogen bonds between residues from different *β*-strands, which are usually not close along the chain [25]. They thus harbour greater geometric variability than *α*-helices and display greater deviations from ideal backbone bond angles [26, 27]. Secondly, *β*-fold membrane proteins mainly have channel or porin conformations, through which molecules can cross the membrane. The internal faces of these *β*-barrels are therefore more hydrophilic even though they are in TM regions. They thus often correspond to weaknesses when computed in lipid environments. Finally, the asymmetrical bilayer of phospholipids in which *β*-barrel proteins are usually inserted have different characteristics than standard phospholipid bilayers. This implies that our statistical potentials generated from TM regions of the whole set of membrane proteins could be less accurate for this kind of proteins (see [3] for further details).

### 3.4 Protein embedding in the membrane

The MPr score can be used not only to identify weak and strong regions but also to predict the protein embedding in the lipid bilayer membrane and more specifically, whether residues are inside or outside the membrane. For that purpose, we transformed the per-residue MPr scores into a binary function, where 0 and 1 mean EM and TM, respectively, using an appropriate threshold value as defined in the Methods section 2.6. We computed the BACC score (Eq. (9)) between the so predicted membrane assignments and OPM annotations, which was found to be equal to 0.88 in a strict protein-level cross validation (section 2.5). This score is very high and we did not expect better as weak residues in TM and EM domains are by definition wrongly predicted, even though they are essential for membrane protein functioning as we will see in the next two subsections.

Note that we used annotations of OPM [16] to train this predictor, and that these annotations are actually also computational predictions. Some discrepancies between the results of our prediction and OPM could therefore be due to errors of OPM rather than of our predictor. We could thus use our predictor to yield new EM/TM assignments, which would differ from OPM assignments in some places. Alternatively, the MPr score could be used as a feature, in combination with other features, to improve the accuracy of the membrane protein embedding in the membrane [16, 28, 29].

### 3.5 Application to Leu transporter

In general, membrane proteins undergo conformational changes to accomplish their biological functions [30, 31]. Residues that are at the basis of these large-scale movements are often stability weaknesses [12] or in other words, frustrated [13]. Indeed, functional constraints prevent them to adopt conformations with high stability contributions. Here we show how we can use the MPr score to gain insight into the functional role of these residues. Note that, to our knowledge, no other method employs dedicated energy functions to study frustration in membrane proteins.

As an example, we analyzed leucine transporter (LeuT), a bacterial homodimeric protein containing twelve transmembrane helices, which uses the electrochemical potential of sodium ions to transport leucine from outside to inside the cell [32]. Both the substrate-free outward-open LeuT structure and the inward-open apo conformation have been resolved via X-ray crystallography [33] (PDB codes 3tt1 and 3tt3, respectively).

We analyzed the difference in membrane propensity index (ΔMpr) between the two conformations in Fig 4.a. We can easily identify the two regions with highest ΔMPr in absolute value: the transmembrane helix 1 (TM1) and the extracellular helix 4 (EL4), which are key elements that allow the opening and closing of the intracellular and extracellular gates, respectively [33]. We thus focus on these two regions and analyze their MPr values in both outward and inward conformations (Fig. 4.b-e).

**Figure 4:**
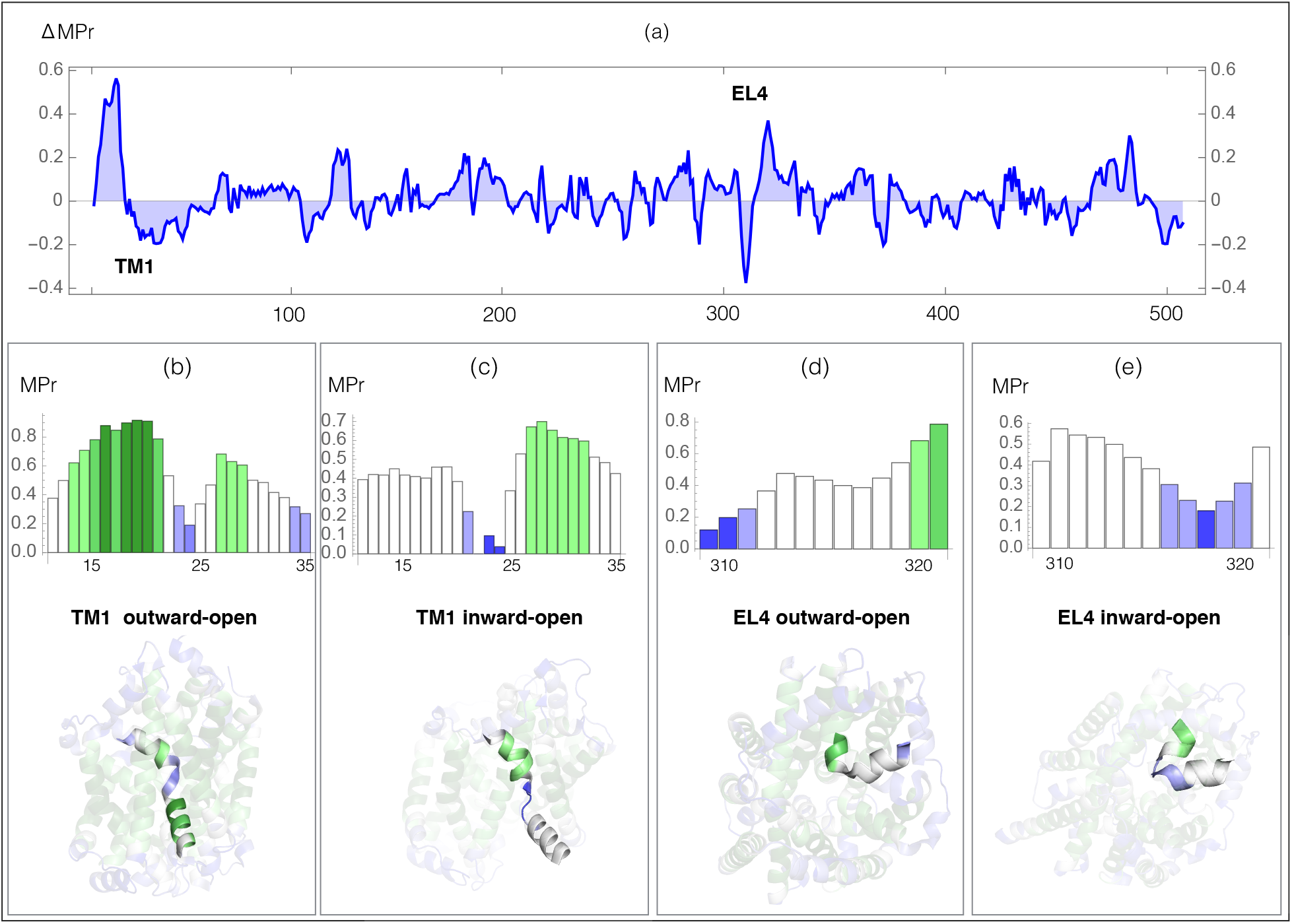
Leu transporter: MPr score in the outward-open (PDB code 3tt1) and inward-open (PDB code 3tt3) conformations (a) Difference in MPr score (ΔMpr) between the two conformations as a function of the sequence position; the positions of the helices TM1 and EL4 are indicated. (b-e) MPr of the TM1 and EL4 helices as a function of the sequence position, and TM1 and EL4 helices embedded in the 3D structure, colored according to the code defined in Table 2; (b) TM1, outward-open conformation; (c) TM1, inward-open conformation; (d) EL4, outward-open conformation; (d) EL4, inward-open conformation.

The transmembrane segment TM1 (residues 11-35) is a kinked helix formed by two helical regions, TM1a (residues 11-21) and TM1b (residues 25-35), separated by a hinge fragment (residues 22-24). TM1a undergoes a large movement upon opening the intracellular gate (Fig. 4.b-c). Despite the fact that the whole TM1 is inside the lipid membrane, the hinge residues have a very low MPr score in both conformations. They are thus stability weaknesses, which is related to their functional role in facilitating the opening and closing of the gate, as confirmed by experiments [33]. The MPr of TM1a changes significantly during the conformational change: in the inward-open conformation, when the intracellular gate is open, TM1 is in only weakly stable (Fig. 4.c), whereas in the outward-open conformation, when the intracellular gate is closed, TM1a establishes stabilizing contacts with TM5 and TM7 which is reflected by high MPr values (Fig. 4.b).

In a similar way, we identified the strong and weak residues in the outer membrane environment related to the conformational movement that EL4 undergoes to open and close the extracellular gate. EL4 (residues 309-321) is a helix of which the last residues (319-321) are unwound and called hinge; its position appears displaced in the outward-open and inward-open conformations. In the outward-open conformation, when the extracellular gate is open, the N-terminal EL4 residues 309-311 are stable in the aqueous environment but the hinge residues are weak (Fig. 4.d). In closing the gate, EL4 changes stability profile, with its N-terminus that becomes weak and its C-terminus and hinge that get more stable (Fig. 4.c) due to extensive contacts including hydrophobic interactions and a hydrogen bond between Ala 319 and Asp 401 in TM10.

This example illustrates how we can use the BRANEart webserver to identify in a simple way residues in membrane proteins that are not optimized for their stability but that have functional roles.

### 3.6 Application to human phospholamban

As second test case, we studied the stability of human phospholamban (PLB), a protein anchored into the cardiac sarcoplasmic reticulum membrane, which is essential to myocardial contractility [34]. Let us start with the analysis of the MPr index as a function of the sequence, computed from the standard pentameric form of PLB (PDB code 1ZLL). As shown in Fig 5.a, the two domains of the protein are easy to identify: the EM part (residues 1-22) has low MPr values and thus the propensity to be stable in water, and the TM part (residues 23-52) has overall much higher MPr values indicating its stability in lipids. A closer look shows that the TM region must be divided in two: the C-terminal part (residues 33-52) is apolar, contains the well known motif LxxIxxx [35] and has high MPr values indicating its stability in lipids, whereas the N-terminus (residues 23-32) is a TM weakness.

**Figure 5:**
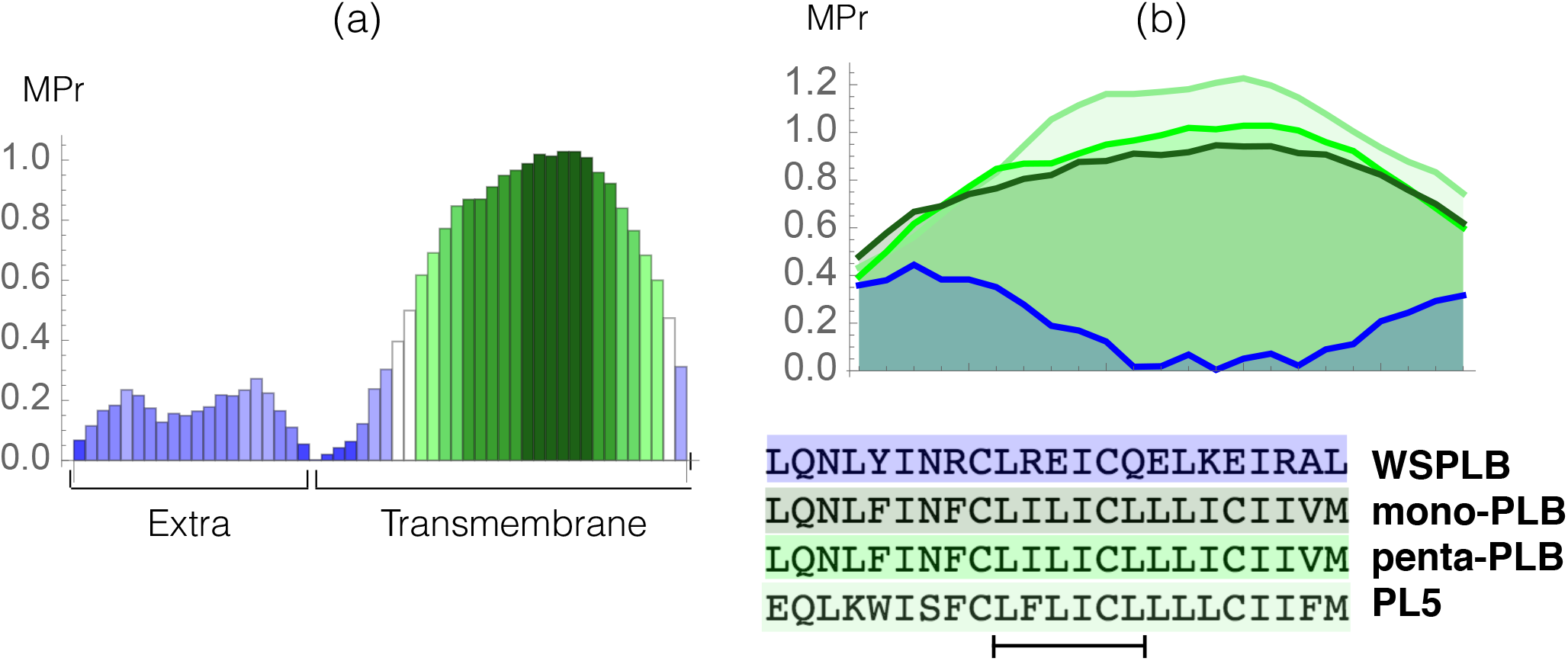
Phospholamban case study. (a) MPr values computed for the pentameric structure of the full-length phospholambam (PDB code 1ZLL); (b) amino acid sequences and MPr values for the TM fragment (residues 28-50) of four different PLB structures: PLB monomer (light green, PDB code 1ZLL), PLB pentamer (green, PDB code 1ZLL), redesigned pentamer PL5 (dark green, PDB code 6MQU) and water soluble PLB tetramer (blue, PDB code 1YOD).

Note that the latter part connects the EM and TM regions and is very close to the water-membrane interface. It thus probably changes dynamically from the lipid TM to the aqueous EM environment and *vice versa*. This prediction is in perfect agreement with both NMR data [36], molecular dynamics simulations [37, 35] and mutagenesis data [38, 39], which identified this region as highly dynamical, unimportant for the stabilization of the structure, but crucial for modulating functional interactions of PLB.

We also analyzed the MPr index of the different types of PLB structures. More specifically, we compared the stability of the wild-type monomeric and pentameric forms (PDB code 1ZLL) [36], of a highly stable designed pentameric variant (PL5, PDB code 6MQU) [35] and of a water-soluble tetrameric variant (WSPLB, PDB code 1YOD) [40]. The MPr values of the TM region (residues 28-50) in all these structures are reported in Fig. 5.b. The TM region of the monomeric form shows a clear preference for the lipid environment with all residues having an MPr value bigger than ∼ 0.5. The pentamer form is predicted to be more stable than the monomer form since inter-chain interactions between the TM fragments strongly stabilize this structural assembly. Note that both the monomer and pentamer forms do exist in phospholipid bilayers even though the latter is dominant. Such oligomer equilibrium is dynamic and can change due to mutations or phosphorylation [41].

The PL5 variant has the largest MPr values and is the most stable form in the lipid environment. This is again in perfect agreement with experiments; in redesigning this protein fragment, the polar residues have been substituted by apolar residues to get stronger interchain interactions driven by apolar sidechain packing and an increased stability of the overall fold [35]. Finally, the redesigned, truncated, water-soluble variant has extremely low propensities to be in a lipid environment, as can easily be seen from our predictions.

### 3.7 BRANEart webserver: features and functionalities

We implemented our structure-based prediction method into the easy-to-use [BRANEart](http://babylone.3bio.ulb.ac.be/BRANEart/help.php) webserver. To run a query, the user first chooses the structure of a membrane protein by either providing its 4-letter code which is then automatically retrieved if available in the PDB, or by uploading a protein structure file in PDB format. The user is then asked to select the relevant chain(s) on which BRANEart has to perform the predictions, taking into account the other chains present in the structure file. Finally, the computation starts.

The main BRANEart output is the MPr score for each amino acid residue in the selected chain(s) of the target protein structure. In addition to returning these values as a downloadable text file, the webserver also displays a table with the MPr of each residue of the targeted protein chain(s), colored according to the code defined in Table 2. In addition, BRANEart provides a multi-featured browser tool to visualize the protein 3D structure, where each residues is colored according to its MPr score. Additionally, there are several advanced visualization functionalities:

- Possibility to hide/show specific chain(s) in the displayed structure, and zoom in and out by left dragging the mouse.
- Possibility to switch to a full screen visualization mode and take a “.png” snapshot of the visualized structures.
- Possibility to select a residue of interest by double clicking on it. The neighboring residues are then displayed in “ball and sticks”.
- Possibility to download a PyMol session file (.pml) to switch to the PyMol representation of the 3D structure.

For further technical details and information about the webserver, we refer to the [BRANEart help page](http://babylone.3bio.ulb.ac.be/BRANEart/help.php).

## 4 CONCLUSION

We presented BRANEart, a computational method to identify the stability strengths and weaknesses in membrane proteins. Extending and combining the newly developed membrane statistical potentials introduced in [3], we defined a MPr score that quantifies whether a residue is stable in a lipid or aqueous environment. Large-scale predictions and applications to test cases show BRANEart’s ability to correctly identify regions in their respective environment that strongly contribute to the stabilization of their host protein, and residues that have instead low impact on stabilization but have functional roles.

Note that our approach can be extended to identify strengths and weaknesses in other environments than the membrane, for example in hot and cold environments using temperature-dependent potentials [21].

We additionally provided a user friendly webserver that, on the basis of the 3D structure of the target membrane protein, computes the MPr score for each residue in the input structure. Visualization tools are provided which simplify the understanding and interpretation of BRANEart results.

To our knowledge, BRANEart is the first accessible, fast and accurate tool that use dedicated membrane protein potentials to identify stability strengths and weaknesses in membrane proteins. The use of such mean force potentials drastically simplifies the analysis of membrane protein stability, since the effect of the lipid environment is implicitly considered. BRANEart can be used in a wide series of applications that range from the analysis of the conformational changes in membrane proteins, the localization of proteins with respect to the lipid membrane, to the identification of residues to target for the rational modification of biophysical characteristics of membrane proteins.

## Acknowledgments

SB wants to convey his sincere gratitude to Dr. Abhirup Bandyopadhyay, Institute De Neurosciences Des Systems, Aix-Marseille University, France for his constant motivation and helpful discussions during the work.

## Data Availability Statement

The list of all the proteins as well as the membrane dependent statistical potentials used in this study can be found in our GitHub repository github.com/3BioCompBio/BRANEart.

